# Incoherent collective cell chemotaxis underlies organ dysmorphia in a model of branchio-oto-renal syndrome

**DOI:** 10.1101/2021.01.27.428404

**Authors:** Augusto Borges, Filipe Pinto-Teixeira, Indra Wibowo, Hans-Martin Pogoda, Matthias Hammerschmidt, Koichi Kawakami, Jerónimo R. Miranda-Rodríguez, Hernán López-Schier

## Abstract

Mutations in the transcriptional co-activator Eya1 cause branchio-oto-renal syndrome (BOR) in humans, and the equivalent conditions in vertebrates. BOR has an incidence of 1/40,000 and is characterized by congenital branchial fistulas, malformations of the inner ear and kidney hypoplasia. Therapeutic interventions for BOR are currently limited to reparative surgery, hearing aids and dialysis. Here we use the mechanosensory lateral line in zebrafish to better understand the role of Eya1 in organogenesis. The lateral line develops from a primordium formed by approximately 150 cells that move together from head to tail of the embryo at a constant velocity. This invariant migration occurs over a trail of Sdf1a chemokine and is controlled by the simultaneous action of two receptors. The CXCR4b is expressed in the front half of the primordium where it acts as a chemokine sensor, whereas the CXCR7b is present in the rear half, serving as a chemokine sink to ensure persistent directionality. We show that the loss of Eya1 strongly reduces the expression of CXCR7b, disrupting the coherent motion of the primordium and leading to lateral-line truncations. We also find evidence of reduced epithelial maturation in primordia lacking Eya1. These findings argue for abnormal collective cell chemotaxis as the origin of organ dysmorphia in BOR.

## INTRODUCTION

The coordinated action of multiple cells governs the development of tissue shape and pattern. For example, collective cell migration influences the formation of various organs, including the eye, kidney, skin follicles, inner ear and lateral line (Scarpa and Mayor, 2016; Dalle Nogare and Chitnis, 2017; Gunawan et al., 2019; La Porta and Zapperi, 2019). The movement of cellular collectives is regulated by a network of cellular communication molecules that include attractive signals, for instance chemokines, and their receptors. One example of collective cell migration that has been extensively studied occurs during the formation of the posterior lateral line in zebrafish (López-Schier, 2010). The lateral line develops from a group of around 150 cells that form a primordium in the embryo (Ghysen and Dambly-Chaudière, 2007). These cells migrate together along the anteroposterior axis of the animal over the course of around 24 hours. As the primordium moves, it deposits clusters of 20–30 cells at semi-regular intervals, which eventually arrest movement and develop into discrete organs called neuromasts. The coherent migration of the primordium - direction, orientation and persistence - is mediated by the complementary expression pattern of the CXCR4b and CXCR7b chemokine receptors (Valentin et al., 2007; Lau et al., 2020). Leading primordial cells express CXCR4b whereas trailing cells express CXCR7b. Both receptors detect the chemokine Sdf1a/CXCL12 that is expressed as a straight line from head to tail along the horizontal myoseptum of the embryo (Donà et al., 2013; Venkiteswaran et al., 2013). The expression of CXCR4b and CXCR7b and epitheliogenesis are under control of intercellular signals acting on non-overlapping parts of the cellular collective. The leading region of the primordium activates Wnt/beta-catenin, whereas FGF signaling is active at the trailing region (Lecaudey et al., 2008; Nechiporuk and Raible, 2008; Aman et al., 2011; Agarwala et al., 2015). Moreover, elegant experiments have revealed that the coherent migration of the primordium requires a combined function of E- and N-cadherins (Colak-Champollion et al., 2019).

Mutations in genes driving collective cell behavior have profound deleterious effects on organogenesis. One gene of particular interest that is co-expressed with CXCR4b and CXCR7b in the lateral-line primordium is Eya1, which when mutated disrupts the distribution of neuromasts (Kozlowski et al., 2005; Sahly et al., 1999; Seleit et al., 2017; Almasoudi and Schlosser, 2021). Eya1 is a protein tyrosine phosphatase with transcriptional activity when associated with the DNA-binding protein Six1. Their function is conserved across species (Shah et al., 2020). For instance, mice lacking Eya1 or Six1 have malformed kidneys and ears (Wong et al., 2013; Xu et al., 1999; Xu et al., 2003; Xu, 2013). In humans, mutations in Eya1 segregate with 40% of cases of Branchio-Oto-Renal syndrome (BOR) (Abdelhak et al., 1997; Sánchez-Valle et al., 2010; Krug et al., 2011). BOR is a heterogeneous condition causing various degrees of renal dysfunction and conductive hearing loss. It characterized by congenital branchial fistulas, malformations of the inner ear and kidney hypoplasia (Kochhar et al., 2007; Feng et al., 2021; Chen et al., 1995). Standard treatments for BOR over the past 25 years have been kidney transplants, dialysis and hearing aids (Smith, 1993; Tian et al., 2022). More innovative interventions are lacking in part because the cellular mechanisms that are disrupted in BOR remain obscure (Soni et al., 2021). Here we combine forward- and reverse-genetic analyses with live imaging to study a model of BOR in zebrafish. Our results shed new light on the role of Eya1 on collective cell migration, and suggest potential avenues to explore novel therapeutic strategies for human patients.

## RESULTS

### Loss of Eya1 disrupts the pattern of neuromast deposition

Alterations of the Sdf1a chemokine or its receptors CXCR4b and CXCR7b lead to defects in neuromast deposition. Therefore, we speculated that genetic mutations affecting the number of neuromasts will identify factors involved in chemokine signaling. Following this rationale, we analyzed zebrafish carrying a loss-of-function mutation in Eya1, which has previously been reported to reduce the number of posterior neuromasts (Fig. 1A-B,E-F) (Kozlowski et al., 2005; Nica et al., 2006). Using somatic CRISPR/Cas9-mediated genome engineering we mutated Eya1 and confirmed that its loss generates an average of 5 neuromasts instead of the 8 organs normally found in wild-type fish at 3 days-post-fertilization (dpf) (Fig. 1C,F). We found a qualitatively similar phenotype in specimens injected with a translation-blocking morpholino against Eya1 (Fig. 1D). To assess the phemotype in more detail, we used two fluorescent enhancer-trap lines: SqET20 to mark supporting cells and SqET4 to highlight mechanosensory hair cells (Parinov et al., 2004). We found that neuromast survival over the course of 4 days after deposition was not affected by the loss of Eya1 (Fig. 1G-J). However, mutant neuromasts had noticeably fewer hair cells (Fig. 1H,J). These data indicate that the lateral-line defects in Eya1 mutants must arise during development and not from postembryonic degeneration of neuromasts. Therefore, we focused attention on the initial formation of the lateral line. When looking at early embryos, we found that loss of Eya1 delayed the migration of primordium (Fig. 2A-E). Some previous work have shown that defective mitotic activity of primordial cells decreases the rate of migration and neuromast deposition (Aman et al., 2011; Gamba et al., 2010; Valdivia et al., 2011; McGraw et al., 2011). However, others have concluded that mitotic activity has negligible effects on migration (Matsuda et al., 2013). Therefore, we assessed cellular proliferation by performing BrdU incorporation in wild-type Eya1(-) embryos. This experiment showed that primordial cells actively divided in both samples (Fig. F-I). Also, because loss of Eya1 decreases cellular viability in several organ primordia, we assessed apoptosis in wild-type and Eya1-mutant lateral-line primordia by the TUNEL assay. We found that whereas most primordial cells in wild-type animals remained viable during migration (Fig. 2J-L), Eya1 mutants experienced significant apoptosis throughout the entire primordium (Fig. M-O). Together, these results show that Eya1 is essential for the initial development of the posterior lateral line.

**Figure 1.**
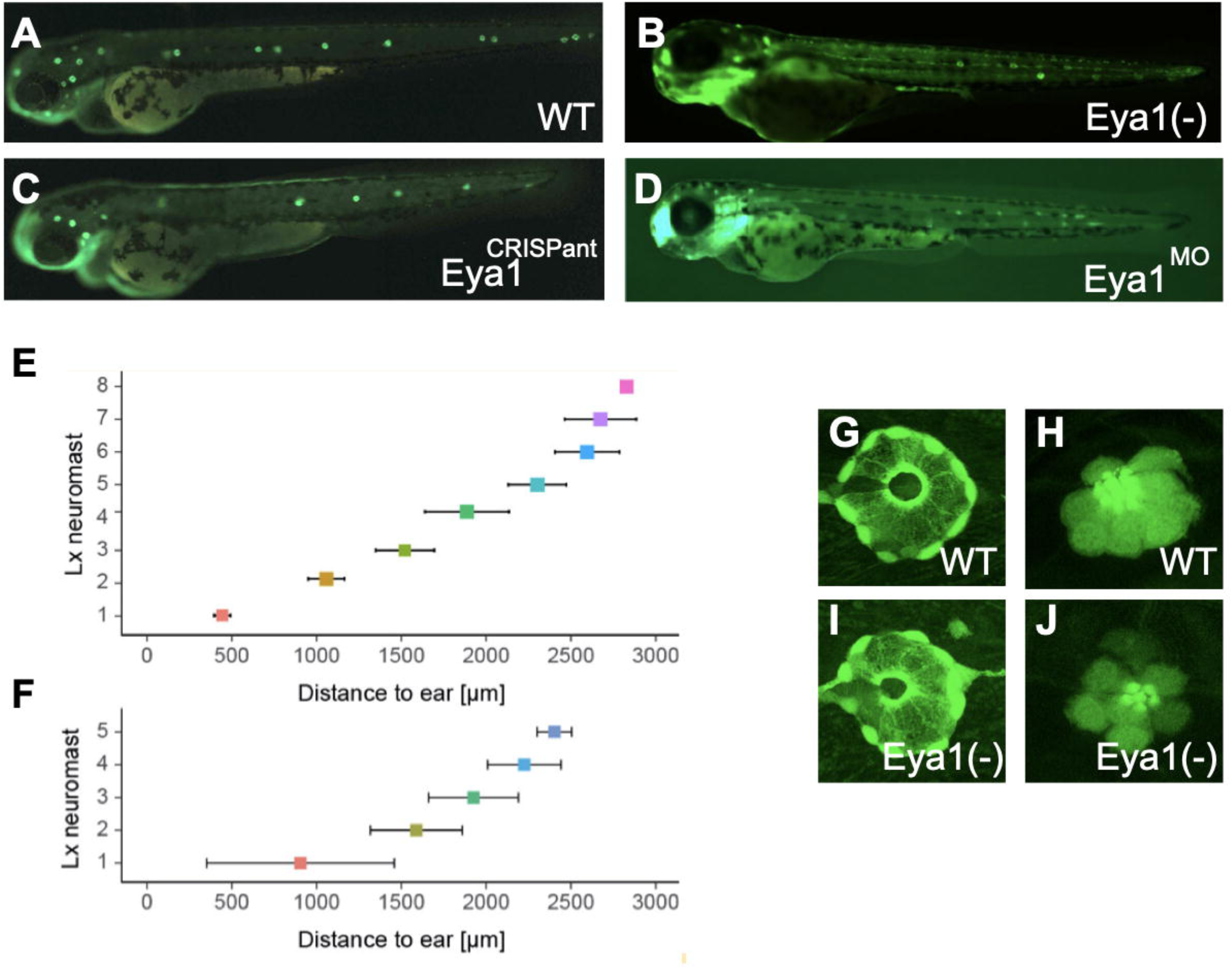
Loss of Eya1 disrupts lateral-line development. **(A-D)** Live images of 72hpf larvae in wild-type (**A**), Eya1 mutant (**B**), Eya1 CRISPant (**C**) and Eya1 morphant (**D**). B-D show qualittatively similar mutant phenotypes in the ouline of the posterior lateral line. **(E-F)** Plot of the distance (in μm) between the caudal limit of the otic vesicle and the average number of deposited neuromasts in wild-type (**E**) and Eya1*-*mutant (**F**) specimens at 3 dpf (mean±s.d.). N= 4 for wild type and N=9 for Eya1-/-. **(G-J)** Live images of neuromasts at 6dpf in wild-type (**G,H**), Eya1 mutants (**I,J**) revealing supprting cells (**G,I**) and hair cells (**H,J**).

**Figure 2.**
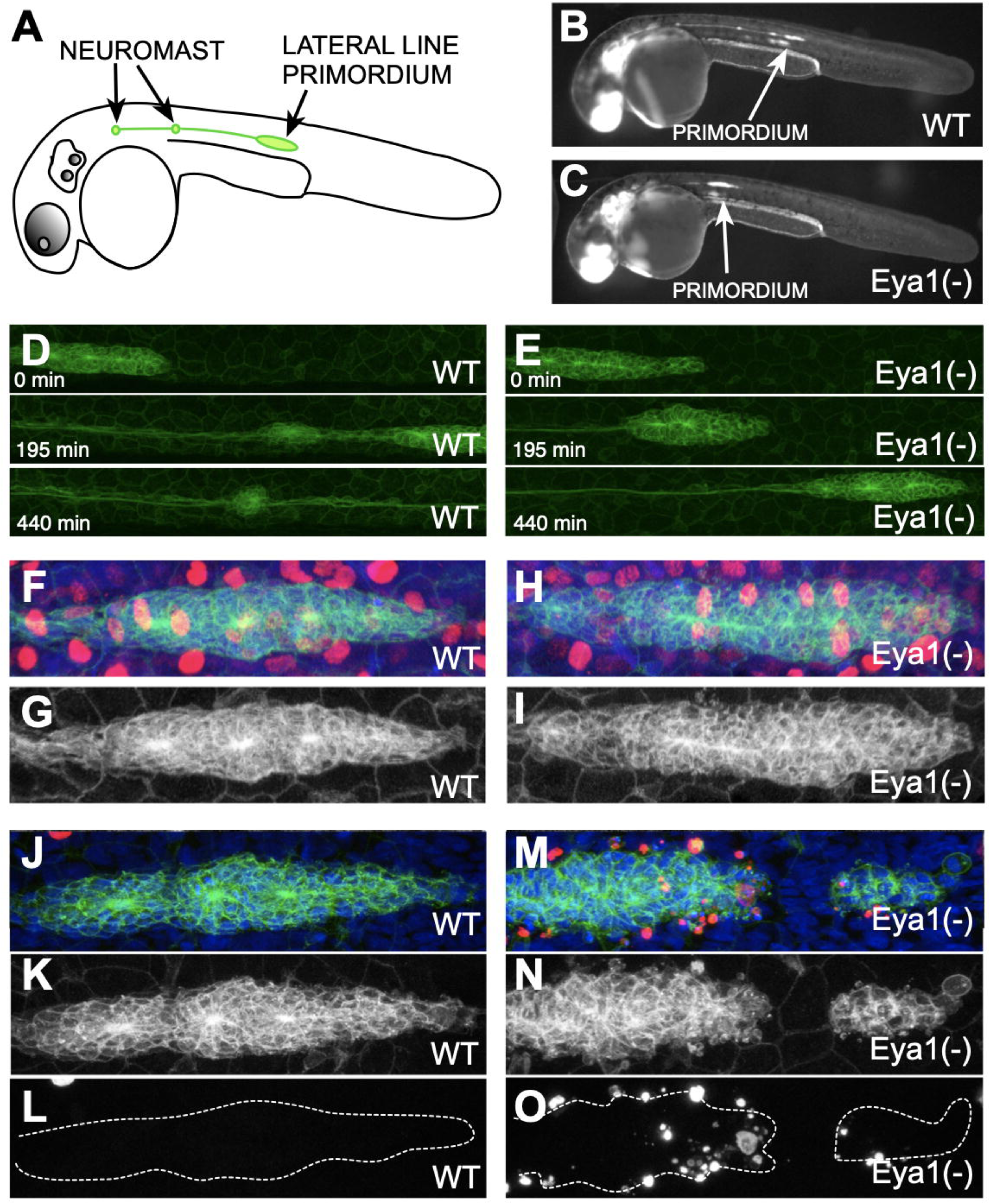
Loss of Eya1 affects the migration and survival of primordial cells. **(A)** Scheme of the developing lateral line in a zebrafish emrbyo. Early neuromasts and the moving primordium are shown in green. **(B-C)** Low magnification confocal images of a wild-type Tg[CldB:lynEGFP] embryo (**B**) and a Eya1-/-;Tg[CldB:lynEGFP] (**C**) at 32 hpf showing the delayed migration of the primordium (white arrows). **(D-E)** High magnification confocal images of a wild type (**D**) and mutant (**E**) primordium. At 195 minutes of migration, the wild-type primordium has deposited one pro-neuromast, whereas the Eya1(-) primordium failed to do so even after 440 minutes. **(F-I)** Cell proliferation of wild-type (**F-G**) and Eya1(-) primordium (**H-I**). BrdU incorporation (red) on a Tg[CldB:lynEGFP] embryos (green in F,H and grey in G-I) shows no evident differences in S-phase completion between representative samples. **(J-O)** Cell viability analysis by the TUNEL assay (red) on Tg[CldB:lynEGFP] embryos (green in J,M and grey in K,N) counterstained with the nuclear dye DAPI (blue) of wild-type (**J-L**) and Eya1-mutant primordia (**M-O**). TUNEL signal alone is shown in L and O. The primordia are outlined by a dotten white line. Notice primrdium split in the mutanrt (**O**).

### Eya1 is necessary for the migratory coherence

Next, we decided to focus on primordial-cell dynamics from the onset of migration by *in toto* videomicroscopy, combining two fluorescent enhancer-trap lines from the Kawakami collection (Kawakami et al., 2004). We first characterized a new line called Tg[gSAG181A], which is unique in that it is the first that expresses EGFP exclusively in the posterior lateral-line primordium (Fig. 3A). Tg[gSAG181A] is an insertion near the SAM and SH3 domain containing 1a (sash1a) *locus* on chromosome 20. The Tg[SAGFF(LF)19A] is an insertion of a Gal4 transgene into Ebf3 *locus* (Kuriki et al., 2020). When combined with a UAS-driven RFP, we observed that it is expressed in the rear part of the primordium and in neuromasts (Fig. 3B). When combined, Tg[gSAG181A;SAGFF(LF)19A;UAS:RFP] allows a specific labeling of the posterior lateral-line primordium for high-quality live imaging and kymography (Fig. 3C-F and Supp. Movies 1 and 2). At a recording temperature of 28°C, we saw that wild-type primordia move at a constant velocity of around 80 μm/hour (Fig. 3C,E), depositing SAGFF(LF)19A-rich neuromasts, whereas Eya1-deficient primordia undergo cycles of migration and stalling, averaging a markedly reduced speed of 14 μm/hour (Fig. 3D,F). The loss of Eya1 strongly reduced the expression of SAGFF(LF)19A (Fig. 3D,F). Incidentally, live imaging of mutant fish shows apoptosis during migration (Supp. Movie 2), supporting our previous result using TUNEL staining (Fig. 2G-G’). Moreover, we found some instances in which the posterior primordium lacking Eya1 loses directionality when it makes u-turns to move back towards the head (Fig. 3F and Supp. Movie 2). Therefore, the loss of Eya1 does not block primordium migration, but instead creates pronounced defects in its otherwise coherent motion.

**Figure 3.**
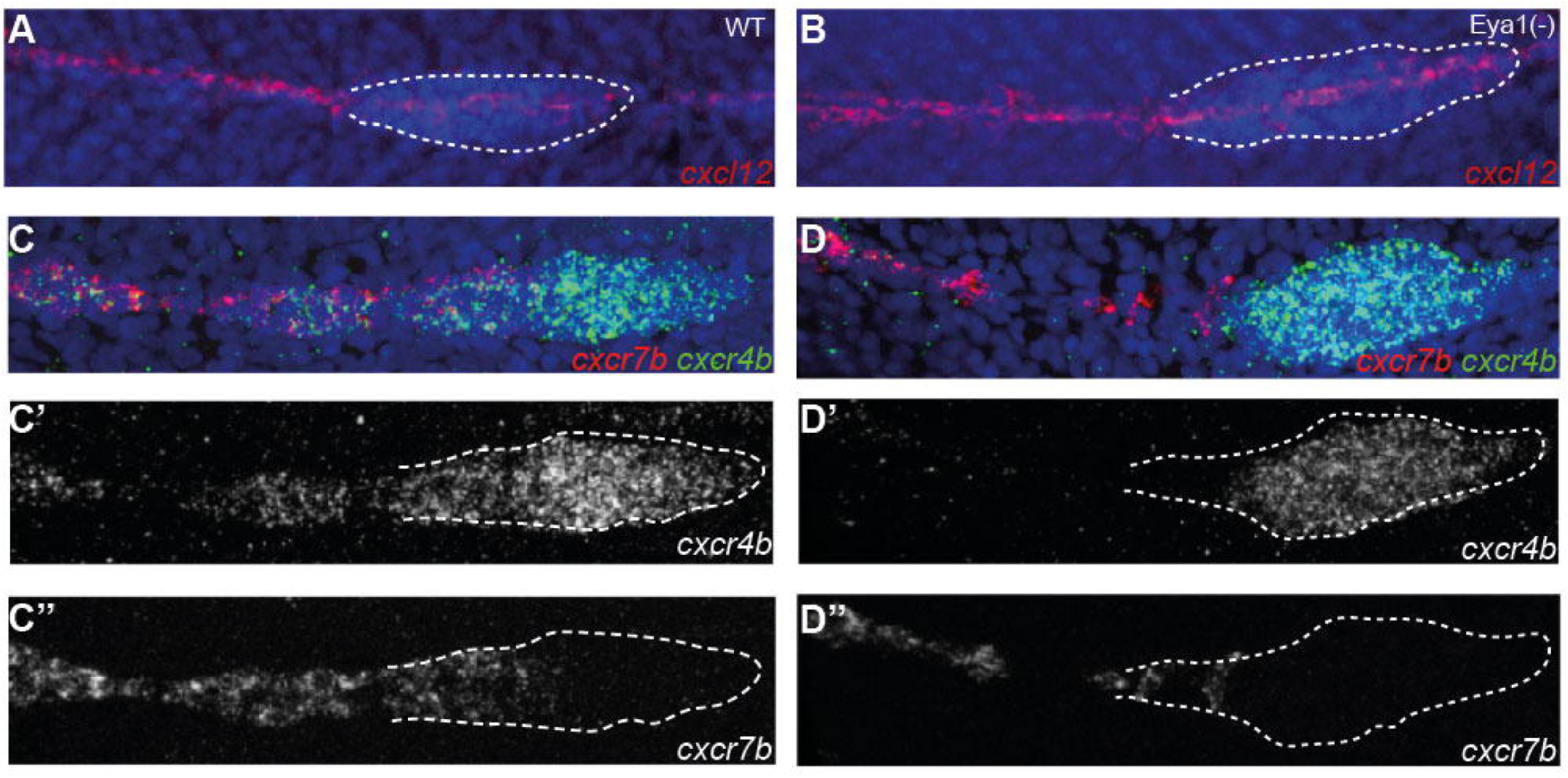
Primordium migration defects in Eya1 mutants. **(A)** Low resolution image of a Tg[gSAG181A] transgenic larva at 30hpf. It specifically expresses EGFP in the posterior lateral line primordium. **(B)** High reolution image of a primordium and a recently deposited neuromast in the Tg[SAGFF(LF)19A;UAS:RFP] transgenic line countertstained with DAPI (blue). SAGFF(LF)19A expresses Gal4 (red) in rear primordial cells, pro-neuromasts and interneuromast cells. **(C)** Still frames of a time-lapse movie focusing on the migrating primordium. RFP driven by SAGFF(LF)19A marks the cells that form rosettes and eventually the proto-neuromast. A few cells expressing 19A:RFP remain in the trailing edge after pro-neuromast deposition. **(D)** Still frames from a representative Eya1 crispant larva in the 181A;19A:RFP transgenic background. No neuromast deposition is seen for the duration of the movie. The fish was confirmed to express 19A:RFP in older neuromasts but barely any red fluorescence is seen in the trailing edge of the primordium. **(E)** Kymograph of the wild-type movie (in C) representing the kinetics of the migration process. The leading edge advances linearly with a velocity of roughly 80 microns per hour at 28°C. The trailing 19A:RFP signal is associated with pro-neuromast rosettogenesis. **(F)** Kymograph of the migration process (in D). The curved shape of the trajectory indicates that after stalling, the primordium performs a u-turn and starts backward migration. Black gaps in the solid green band reflect transient splitting events of the primordium. **(G)** Stillframes of the leading edge in a primordium lacking Eya1 and observed in a Tg[CldB:lynEGFP] transgenics. From 20’ to 50’, a handful of leading cells split from the rest of the primordium but subsequently fuse back.

### Eya1 is necessary for CXCR7b expression

Using the Tg[Cldnb:lynEGFP] transgenic line to individualize cells, we also found many instance in which the primordium splits transiently during migration (Fig. 3G). Interestingly, both phenotypes have been observed in fish lacking CXCR7b (Valentin et al., 2007). Therefore, we decided to analyze the signaling pathway that governs the collective chemotaxis of primordial cells: the CXCR4b and CXCR7b chemokine receptors and the Sdf1a chemokine. We hypothesized that loss of Eya1 does not affect Sdf1a because migration in the mutants occurred along the horizontal myoseptum, which is the normal path of the primordium. Indeed, the expression of Sdf1a was normal in Eya1-mutant fish (Fig. 4A-B). By contrast, we saw that while CXCR4b remained expressed in the leading zone in Eya1 mutant primordia (Fig. 4C-C”), the expression of CXCR7b was strongly reduced (Fig. 4D-D”). We conclude that the function of Eya1 is intrinsic to the primordium and that its loss causes incoherent primordium migration due to strongly reduced CXCR7b expression.

**Figure 4.**
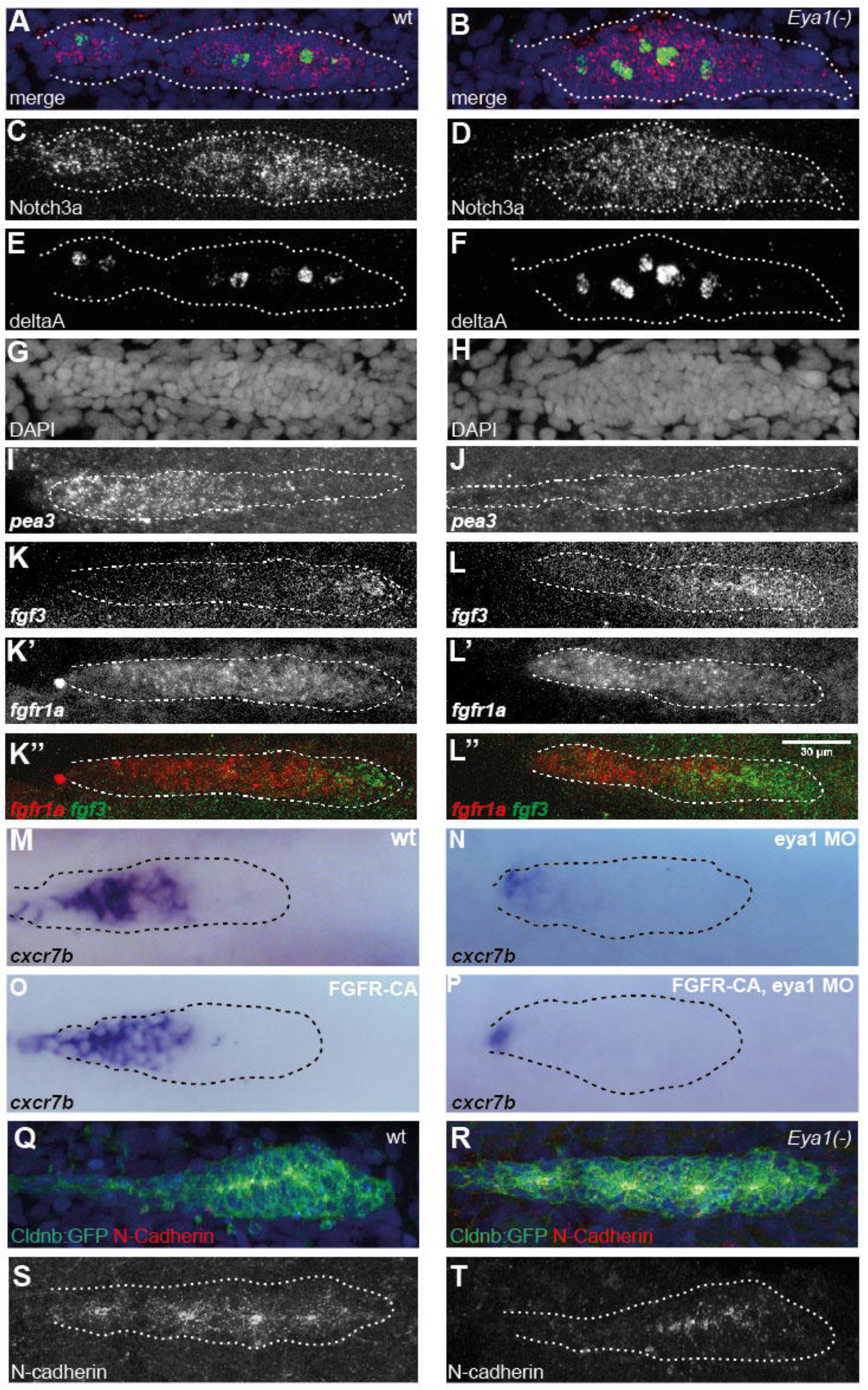
Eya1 is necessary for Cxcr7b expression. **(A,B)** Fluorescent whole-mount *in situ* hybridization of Sdf1a (red), counterstained with DAPI (blue) to reveal the nuclei for better identification of the primordium (white dotted outline). It shows that the Sdf1a gene (red) is expressed along the horizontal myoseptum in the wild type **(A)** and Eya1 mutants **(B)**. **(C-D’’)** CXCR4b (green) and CXCR7b (red) gene-expression profiles in wild type **(C)** and Eya1 mutants **(D)** The CXCR4b gene is strongly expressed in the leading region of primordium in both in wild-type and Eya1 mutants **(C’,D’)**. The expression of CXCR7b, however, is strong in the trailing region of the wild-type primordium (overlapping with CXCR4b), but almost completely lost in Eya1 mutants, as it is restricted to the very end of the trailing region and never overlaps with CXCR4b **(C’’,D’’)**.

### Loss of Eya1 affects FGF signaling and delays epitheliogenesis

Our previous results show that Eya1 controls the expression of a subset of genes expressed in the primordium. To further explore this, we profiled gene expression by whole-mount *in situ* hybridization. We first looked at the Notch pathway because it reveals the rate of cellular differentiation in the primordium (Matsuda et al., 2013). We found that the expression of the Notch3a receptor and the DeltaA ligand were not affected in Eya1 mutants (Fig. 5A-H). Next, we focused on FGF signaling pathway for two reasons: 1) its activity domain overlaps that of CXCR7b and 2) blocking FGF results in migratory defects (Nechiporuk and Raible, 2008; Lecaudey et al., 2008). In fish lacking Eya1, the expression of Pea3, a *bona fide* target of FGF signaling, was almost completely abolished (Fig. 5I-J). Eya1 loss also reduced the expression of Fgfr1a, which is normally maintained by a positive feedback loop in the FGF signaling cascade (Fig. 5K-L’’), but slightly expanded the expression domain of the Fgf3 ligand (Fig. 5K-L’’). Thus, in the absence of Eya1 the primordium experiences simultaneous reduction of FGF signaling and CXCR7b expression. This combination of phenotypes has already been reported for zebrafish embryos treated with the FGFR inhibitor SU5402, suggesting that loss of Eya1 reduces CXCR7b expression by impairing FGF signaling (Aman et al., 2011). To test this possibility, we artificially forced FGF signaling in Eya1 mutants. To this end, we abrogated Eya1 using a translation-blocking morpholino and activated FGF using the transgenic line Tg[hsp:CA-FGFR1] that expresses a constitutively active form of the FGFR1 upon heat-shock (Lee et al., 2009). As expected, knockdown of Eya1 nearly completely eliminates CXCR7b expression (Fig. 5M-N). However, constitutive FGF signaling did not expand the expression of CXCR7b in wild-type fish (Fig. 5O), nor did it rescued CXCR7b expression in animals lacking Eya1 (Fig. 5P). These results indicate that Eya1 induces CXCR7b expression independently or downstream of the FGF receptor. Finally, it has been amply documented that FGF signaling governs epitheliogenesis during the maturation of neuromast in the primordium (Nechiporuk and Raible, 2008; Agarwala et al., 2015; Lecaudey et al., 2008; Kozlovskaja-Gumbrienė et al., 2017; Durdu et al., 2014; Harding et al., 2014; Neelathi et al., 2018). Also, that the epithelial and neuronal cadherins are necessary for the formation of epithelial rosettes that prefigure neuromasts (Revenu et al., 2014). Therefore, we assessed epitheliogenesis by staining wild-type and Eya1-mutant specimens with an antibody to N-cadherin, which highlights the constricted apices of the epithelial cells that form the rosettes. This experiment showed that although epithelialization does occur in the mutants, rosette formation is strongly defective (Fig. 5Q-T). Because neuromasts eventually do form (Fig. 1G-J), we conclude that loss of Eya1 or delays epithelial maturation.

**Figure 5.**
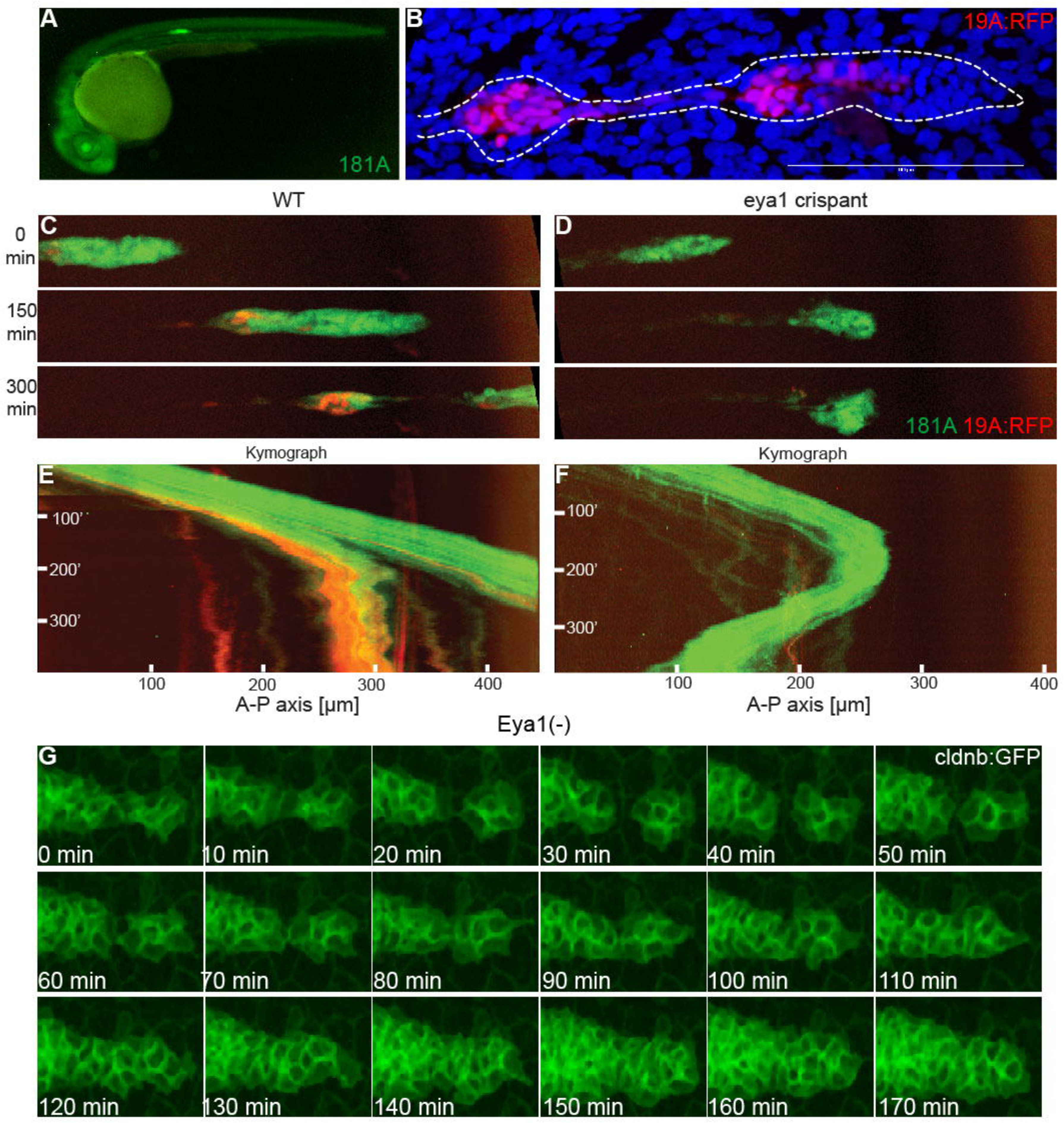
Loss of Eya1 disrupts FGF signaling and epitheliogenesis in the primordium. **(A-H)** Notch pathway in wt and Eya1 (-) primordia. In situ hybridization against Notch3 **(C,D)** and DeltaA **(E,F)**. **(I-L’’)** HCR-FISH in whole-mount larvae for FGF pathway markers pea3 (etv4), fgf3 and fgfr1a. In wild-type primordia, pea3 expression is strongest in the trailing end **(I)**. When Eya1 is somatically eliminated by CRISPR, pea3 is almost completely lost **(J)**. Fgf3 ligand is produced normally in the leading tip of the primordium from which it diffuses posteriorly and binds to the Fgfr1a receptor expressed in trailing cells **(K-K’’)**. Eya1 crispants show a slight expansion of fgf3 distribution while the Fgfr1a trailing domain recedes mildly **(L-L’’)**. **(M-P)** WM-ISH to Cxcr7b in the posterior lateral-line primordium in wild type controls **(M),** Eya1 morphants **(N),** Tg[hsp70:ca-Fgfr1] transgenics **(O)** and Tg[hsp70:ca-Fgfr1] transgenics + Eya1 morphants **(P)**. Overactivation of FGF signaling does not expand the expression domain of Cxcr7b **(O),** and does not rescue the down-regulation of Cxcr7b expression upon loss of Eya1 **(P)**. **(Q-T)** Antibody staining of N-cadherin (red) to reveal adherens junction and the apical constriction of rosettes in Tg[CldB:lynEGFP] embryos (green). Apical constriction of the rosettes are evident in the wild type sample **(S)**, but they are disrupted in Eya1 mutants **(T)**.

## DISCUSSION

The zebrafish has emerged as a powerful model to understand the fundamentals of human disease (Santoriello and Zon, 2012). Here we show that the activity of the Eya1 is essential for the coherent migration of the posterior lateral-line primordium. Quantitative live imaging revealed that Eya1-mutant primordia do not maintain migratory persistence and often undergo fragmentation and directional reversion of movement. Interestingly, this combination of defects has been seen in zebrafish devoid of the chemokine receptor CXCR7b, but not in those lacking CXCR4b or the chemokine Sdf1a (Valentin et al., 2007). Therefore, we reasoned that the Eya1-mutant phenotype derives from defects in CXCR7b. We confirm this prediction by showing that primordia lacking Eya1 failed to express CXCR7b. In addition, loss of Eya1 reduced epithelial maturation. This phenotype has been seen upon pharmacological or genetic inhibition of FGF signaling (Nechiporuk and Raible, 2008; Agarwala et al., 2015; Lecaudey et al., 2008). Indeed, we found evidence of reduced FGF signaling in the primordium of fish lacking Eya1. This conclusion is supported by recent findings showing that FGF signaling controls the expression of CXCR7b in the primordium (Dries et al., 2021). However, our experiments suggest that the effect of Eya1 and FGF signaling upon CXCR7b may be independent or, alternatively, that Eya1 mediates FGF-dependent CXCR7b expression downstream of the activated receptor. Further dissection of how Eya1 controls the chemokine cascade in the zebrafish lateral line will yield novel insights on the fundamental mechanisms that underlie coherent multicellular behavior.

Importantly, our work has clinical significance. Genetic polymorphisms in Eya1 segregate with 40% of BOR cases in humans (Krug et al., 2011; Kochhar et al., 2007; Sánchez-Valle et al., 2010). Yet, the cellular processes affected by mutations in Eya1 and its effectors remain undefined. Collective cell migration underlies morphogenesis of the inner ear and the kidney, the two main organs affected in BOR patients (Vasilyev et al., 2009; Schumacher, 2019; Ishii et al., 2021; Renauld et al., 2022). Based on our findings, we propose that Eya1 governs coherent collective cell movement during otic and renal development mainly via chemokine signaling. However, it is important to note that BOR patients express a wide spectrum of defects, which may be explained by Eya1 controlling multiple processes in parallel, either directly or indirectly, via various molecular pathways. This hypothesis is supported by genome-wide studies in humans, which have identified Fgf3 as a contributor to BOR. It is noteworthy experiments *ex vivo,* which have implicated FGF signaling in ear organogenesis in birds and mammals. For example, depletion of Fgf3 in placodal explants blocked the formation of otic vesicles, and homozygous Fgf3 mutant mice develop severely malformed ears (Represa et al., 1991; Hatch et al., 2007). Thus, our data support the idea that phenotypic variation in Eya1-linked BOR results from coincident loss of FGF signaling and multicellular chemotaxis. Mutations in the obligate Eya1-partner Six1 are also causative of BOR. Notably, CXCR7b is a presumptively direct target of Six1 and Eya1 in *Xenopus* cranial placodes, which include the inner ear and the lateral line (Riddiford and Schlosser, 2016). Given that the causative mutation for over 50% of BOR cases is not yet known, we predict that the lateral line of zebrafish will remain a powerful model to validate genomic polymorphisms from GWAS studies of BOR patients, and generate novel cellular and molecular insights with translational potential. On this regards, our findings raise the possibility that augmenting residual CXCR7 activity may improve the outcome of Eya1 mutations in humans (Jiang et al., 2021; Hughes and Nibbs, 2018). It also encourages the development of tissue engineering approaches to control collective cell migration aimed at clinical applications (Manivannan et al., 2012).

## ACKNOWLEDGEMENTS

We thank K. Poss, G. Weidinger, and T. Whitfield for the gift of zebrafish lines and DNA constructs. This work was funded by the NIH BRAIN Initiative grant 1U19NS104653-01, the BMBF grant 01GQ1904, and discretionary faculty funding from the New York University Abu Dhabi to HL-S. JRM-R received funding from the European Union’s Horizon 2020 research and innovation programme under the Marie Sklodowska-Curie grant agreement No. 840834.

## CONFLICT OF INTEREST STATEMENT

HL-S is scientific advisor and paid consultant for Sensorion (France). The company had no role in this study. No conflict of interests exists.

## MATERIALS AND METHODS

### Zebrafish animals and strains

Fish used were maintained under standardized conditions. Experiments were performed in accordance with protocols approved by the Ethical Committee of Animal Experimentation of the Helmholtz Zentrum München, the German Animal Welfare act Tierschutzgesetz §11, Abs. 1, Nr. 1, Haltungserlaubnis according to the European Union animal welfare, and under protocol number Gz.:55.2-1-54-2532-202-2014 and Gz.:55.2-2532.Vet_02-17-187 from the “Regierung von Oberbayern” (Germany). Eggs were collected from natural spawning and maintained at 28.5°C. Embryos were staged by hours post fertilization (hpf). The Eya mutant allele used in this study is *dog*^*tm90* (Nica et al., 2006). Embryos were genotyped according to Kozlowski et al. (Kozlowski et al., 2005). Transgenic lines used were sqgw57AEt(Pinto-Teixeira et al., 2015), Tg[Cldnb:lynEGFP] (Haas and Gilmour, 2006), Tg[hs70:CA-FGFR1](Lee et al., 2009), gSAG181A and SAGFF(LF)19A (Kawakami et al., 2004). Heat-shock treatments were performed by immersing embryos in the E3 medium in 2-ml tubes in a water bath at 39°C for 20 minutes.

### Translation-blocking antisense morpholinos

cRNAs were synthesized using mMessage mMachine (Ambion) according to manufacturer’s instructions. Morpholinos that were used in this study were: MO1Eya1 (5’- AAACAAAGATGATAGACCTACTTCC-3’ (Kozlowski et al., 2005).

### Somatic CRISPR gene knock-out

Crispr somatic mutagenesis of Eya1 was done with 4 sgRNAs (Wu et al., 2018). A 1 mg/ml equimolar mixture of 4 sgRNAs, transcribed with MEGAshortscript T7 (Thermo Fischer), 5 mM Cas9 protein (Sigma), and 300 mM KCl, was injected into one-cell stage embryos. The sequences in the eya1 gene targeted by each sgRNA are the following: CTTCCACTTACTCGGCTGTG, TTGTCAATGTTGGGACCGTT, GACGTACCTTCAGTGCCATT, AGAGCCGTCTGCTACAGAGG

### BrdU treatment

10 mM 5-Bromo-2’-deoxyuridine (BrdU, B5002, Sigma) stock solution in DMSO was diluted to 10 μM in E3 medium and used to soak embryos. BrdU was incorporated into newly synthesized DNA in the S-phase; therefore, it functioned as a marker for cell proliferation. Embryos at 24 hpf were dechorionated and allowed to develop in this solution until desired stages. Embryos were fixed in 4% Paraformaldehyde (PFA) overnight at 4°C and then used for immunohistochemistry.

### TUNEL assay

Apoptosis in the migrating primordium was identified using terminal transferase-mediated dUTP nick end-labeling (TUNEL) assay according to manufacturer’s instruction with minor modifications (In situ Cell Death Detection Kit, TMR Red, Roche). Embryos at 30-42 hpf were dechorionated and fixed in 4% PFA overnight at 4°C, then stored in 100% methanol at −20°C for at least 1 day. They were rehydrated, permeabilized in 10 μg/ml Proteinase K in 0.1% PBSTw and post-fixed in 4% PFA then washed several times in 0.1% PBSTw. Embryos were then incubated in fresh TUNEL buffer for 1 h followed by incubation in the TUNEL reaction mix for 3 hrs at 37°C in dark. As negative control, embryos were incubated in TUNEL buffer only. Positive control embryos were incubated in polymerase chain reaction buffer containing 3 U/ml DNase I recombinant (Roche) for 1 h at 37°C before adding the TUNEL reaction mix. After the reaction, samples were washed several times in 0.1% PBStw at room temperature and stored in 0.1% PBSTw containing Vectashield mounting medium with DAPI (VectorLabs).

### Whole-mount immunohistochemistry

Staged embryos were dechorionated and fixed in 4% PFA overnight at 4°C and washed several times with 0.1% Tween-20-containing Phosphate Buffer Saline (0.1% PBSTw). Larvae were blocked in 10% Bovine Serum Albumin (BSA) for at least 2 hours. Incubation with primary antibody was done overnight at 4°C. Primary antibodies and monoclonal antibodies were used at the following dilutions: mouse monoclonal antibody anti-BrdU, 1:100 (Upstate), rabbit anti-N-cadherin, 1/500. Texas red-labeled donkey anti-mouse and -rabbit and Cy5-labeled donkey anti-mouse and – rabbit immunoglobulin secondary antibodies (Jackson ImmunoResearch) were used at 1/150. For BrdU labeling detection, before blocking and applying primary antibody anti-mouse, additional steps were needed to permeabilize the nuclear membrane and to denature DNA strands. Larvae were incubated with 10 μg/ml Proteinase K in 0.1% PBSTw for 20 min at room temperature. Immediately afterwards, larvae were post-fixed in 4% PFA for 15 min and washed several times in 0.1% PBSTw. To denature DNA, samples were incubated in fresh 2 N HCl for 1 h and washed several times in 0.1% PBSTw. Samples were blocked in BrdU blocking solution for at least 2 hrs at RT and then incubated in BrdU blocking solution containing the antibodies at 4°C.

### Whole-mount *in situ* hybridization

For ISH, antisense digoxigenin- and fluorescein-labeled riboprobes were synthesized according to manufacturer’s instructions (Roche) by using T7/SP6/T3 RNA polymerases. Probes used were: Sdf1a, Cxcr7b, Cxcr4b, Pea3. Whole-mount two-color fluorescence ISH was performed using anti-DIG and -fluorescein POD antibodies (Roche) and Tyramide Signal Amplification (TSA, PerkinElmer) to detect the riboprobes. Larvae were mounted in 0.1% PBSTw with Vectashield/DAPI (1/100, Vector Labs.) For HCR whole-mount fluorescent *in situ* hybridization, a set of 20 probe pairs was used (Molecular Instruments, Inc.) following a protocol described by the manufacturer. Briefly, samples were fixed in 4% paraformaldehyde (PFA) for 24h at 4°C, permeabilized with methanol and cooled to 20°C. Next day, samples were rehydrated, treatment with proteinase K and post-fixed in PFA for 20 min at room temperature. The samples were washed with PBST between the steps. Probe hybridization buffer was used for the prehybridization for 30 min at 37°C and the samples were incubated in the probe solution, prepared following the manufacturer’s instructions, overnight at 37°C. After removing the probe solution, washing the samples and incubating them in the pre-amplification buffer, the samples were incubated in the hairpin mixture overnight in the dark at room temperature. Finally, after several washes with SSCT, the cell nuclei were stained with DAPI (40,6-diamidino-2-phenylindole, Sigma) 1 hour at room temperature.

### Imaging and time-lapse video microscopy

For whole-mount ISH, embryos were deyolked, flat mounted and photographed with a Olympus BX61 microscope using 20X or 40X dry objectives with transmission light. Whole embryo images were acquired on a Leica MZ10 stereomicroscope. Fluorescent images were acquired using either a Leica SP5 or SPE microscope using 20X dry objective or 40X oil immersion objective. Images were processed using Imaris and/or ImageJ software packages, and assembled with Adobe Photoshop CS2, Adobe Illustrator CS2, and Macromedia FreeHand MX. For time-lapse imaging, staged and dechorionated embryos were anesthetized with tricaine and mounted in 0.8-1% low-melting-point agarose on a glass-bottom culture dish (MatTek) as previously described (Pinto-Teixeira, F. eat al., 2015). Z-stack series were acquired every 4-10 min using a 20X dry objective of Leica SPE or SP5 confocal microscope. All movies were processed with the Imaris or ImageJ software packages. An unpaired two-tailed T test with Welch’s correction was used to compare the position of neuromast L4 in Eya1 mutants and wild-type siblings. Statistics were performed using the GraphPad Prism software and Excel running QI Macros.

## Notes

### Summary of Updates

The manuscript was revised to correct grammatical and factual errors in the text, as well as to update the affiliation of three authors

